# Automated Continuous LLM-driven Biomedical Literature Mining for a Large-Scale Hepatotoxicity Knowledge Base

**DOI:** 10.64898/2026.11.09.765369

**Authors:** Ji Qiu, Yamini Chitikela, Zhenquan Jia, Chunjiang Zhu

**Affiliations:** Columbia University, New York, NY, USA; Old Dominion University, Norfolk, VA, USA; UNC Greensboro Greensboro, NC, USA

**Keywords:** literature mining, large language models, information extraction, hepatotoxicity, QSAR, graph neural networks

## Abstract

Predicting whether a chemical disturbs liver and mitochondrial function matters for drug safety and environmental risk assessment, but the data available for training such models are small and mostly curated by hand. We describe an automated pipeline that uses a large language model (LLM) to collect hepatotoxicity findings from the biomedical literature. The pipeline runs a PubMed/PMC query, downloads the matching full texts or abstracts, asks the LLM to record each reported effect in a fixed schema (tested chemical, biomarker, direction of change, and related fields) with controlled vocabularies, and writes de-duplicated records to a database that is updated as new papers appear. The current snapshot holds 189,212 findings on 17,002 chemicals from 53,575 articles (1995–2026), covering 235 biomarkers reported for at least 20 chemicals. In a manual check of 217 findings from 35 articles, 208 findings (from 33 articles) were fully correct; the other 9, all from 2 articles, were incorrect. As a use case, we train structure-based QSAR models, both graph neural networks and classic baselines, for each biomarker; at this data scale the classic models remain competitive. The pipeline and the knowledge base will be made publicly available.

## I. INTRODUCTION

Chemical databases built around specific hepatotoxicity endpoints are a basic resource in toxicology. They supply lists of suspect substances, support hazard and risk assessment [1], and provide training data for quantitative structure–activity relationship (QSAR) models that predict the effects of untested chemicals [2]. Public toxicity data are still limited, though. Most come either from high-throughput in vitro screening, such as the Tox21 programme [3], [4] and assays from the Broad Institute and the Scripps Molecular Screening Center [5], or from literature-based databases such as ChEMBL [6] and the Comparative Toxicogenomics Database (CTD) [1]. The literature-based resources depend on manual curation, which is slow and costly and cannot keep up with the growing literature.

The liver is the main site of drug and xenobiotic metabolism, and chemical-induced liver injury is a leading cause of failure in drug development and of the withdrawal of marketed drugs [7]. Hepatotoxicity is usually characterized with biomarkers such as alanine aminotransferase (ALT), aspartate aminotransferase (AST), alkaline phosphatase (ALP), and bilirubin, together with mitochondrial indicators such as reactive oxygen species (ROS), malondialdehyde (MDA), and ATP [8], [9]. Many experimental studies have measured how chemicals change these markers, but their results are scattered across decades of papers. No structured, modeling-ready resource links chemicals to their hepatotoxic effects, which makes it harder to screen chemicals and to rank high-hazard compounds.

Large language models (LLMs) can read free text and return structured information without hand-written rules or annotated training corpora. In this work, we use an LLM to build and evaluate a hepatotoxicity literature-mining workflow. Our contributions are:

- **A large-scale hepatotoxicity dataset**. Each finding records the tested chemical, test subject, biomarker, effect direction, exposure conditions, and a verbatim evidence sentence that links it to its source article.
- **An automated, continuously updated pipeline**. New articles are downloaded, findings are extracted with Gemini 2.5 under a fixed schema and controlled vocabularies, and records are written to a de-duplicated database. A daily job adds new papers without manual work or a database rebuild.
- **QSAR baselines**. For each of the 74 markers with enough data, we train models that predict a chemical’s effect direction (increase, decrease, or no significant change) from structure alone, and we compare three graph neural networks (Graphormer, GraphGPS, and GraphSAGE) with random forest and XGBoost under identical data splits.

## II. RELATED WORK

### LLM-based Information and Relation Extraction

Extracting structured knowledge from scientific text is a long-standing text-mining task. Earlier systems used rules, named entity recognition (NER), and supervised relation extraction. They needed hand-built rules or annotated corpora and mostly handled binary or ternary relations, such as drug–disease or gene–phenotype pairs. LLMs have made extraction more flexible and have been applied to biomedical relation extraction [10]–[12]. Mature resources also exist. PubTator3 [13] automatically annotates entities (e.g., chemicals, genes, and diseases) and selected relations across PubMed and PMC, and CTD [1] curates chemical–gene, chemical–phenotype, and chemical–disease interactions through expert curation supported by text-mining triage. Our work complements these resources. Instead of general entity or relation annotation, we extract experiment-level hepatotoxicity findings that record the chemical, biomarker, test subject, exposure conditions, and effect direction together under controlled vocabularies. To our knowledge, existing tools do not provide this multi-field schema directly.

### Automatic Knowledge Base and Dataset Construction

Several studies have used LLMs to build datasets from papers. In materials science, conversational LLMs with prompt engineering [14] and fine-tuned LLMs for joint NER and relation extraction [15] have been used to compile materials property and chemistry data. In toxicology, GPT-based text mining has produced lists of prooxidants and antioxidants [16], and literature mining combined with LLMs has been used to rank candidate hepatotoxicants at the compound level [17]. These studies show that LLM-based dataset construction is feasible. Our work differs in scope: we extract experiment-level findings for many hepatotoxicity biomarkers rather than one endpoint, process more than fifty thousand articles with a pipeline that keeps updating, and provide QSAR baselines on the result.

### Hepatotoxicity Datasets and QSAR Modeling

Curated references such as the FDA’s DILIrank and the NIH’s LiverTox give authoritative annotations of drug-induced liver injury (DILI), and many QSAR and machine learning models have been trained on them. These resources are small (hundreds to about a thousand drugs) and carry a single binary label (DILI concern or not), so they cannot describe how a chemical affects individual biomarkers. We instead build a large, automatically constructed chemical–effect dataset across many biomarkers and use it to compare graph neural networks (GNNs) with classic models.

## III. METHOD

### A. Overview and Problem Formulation

PubMed is a bibliographic database of the U.S. National Library of Medicine (NLM) that indexes citations and abstracts; PubMed Central (PMC) is NLM’s free full-text archive. Our pipeline has four stages (Fig. 1). Articles are collected from PMC (full text) or PubMed (abstract); Gemini 2.5 extracts findings with a fixed prompt; the findings are stored in a SQLite database that is refreshed daily; and the cleaned data are used to train QSAR models that predict the effect direction (increase, decrease, or no significant change).

**Fig. 1.**
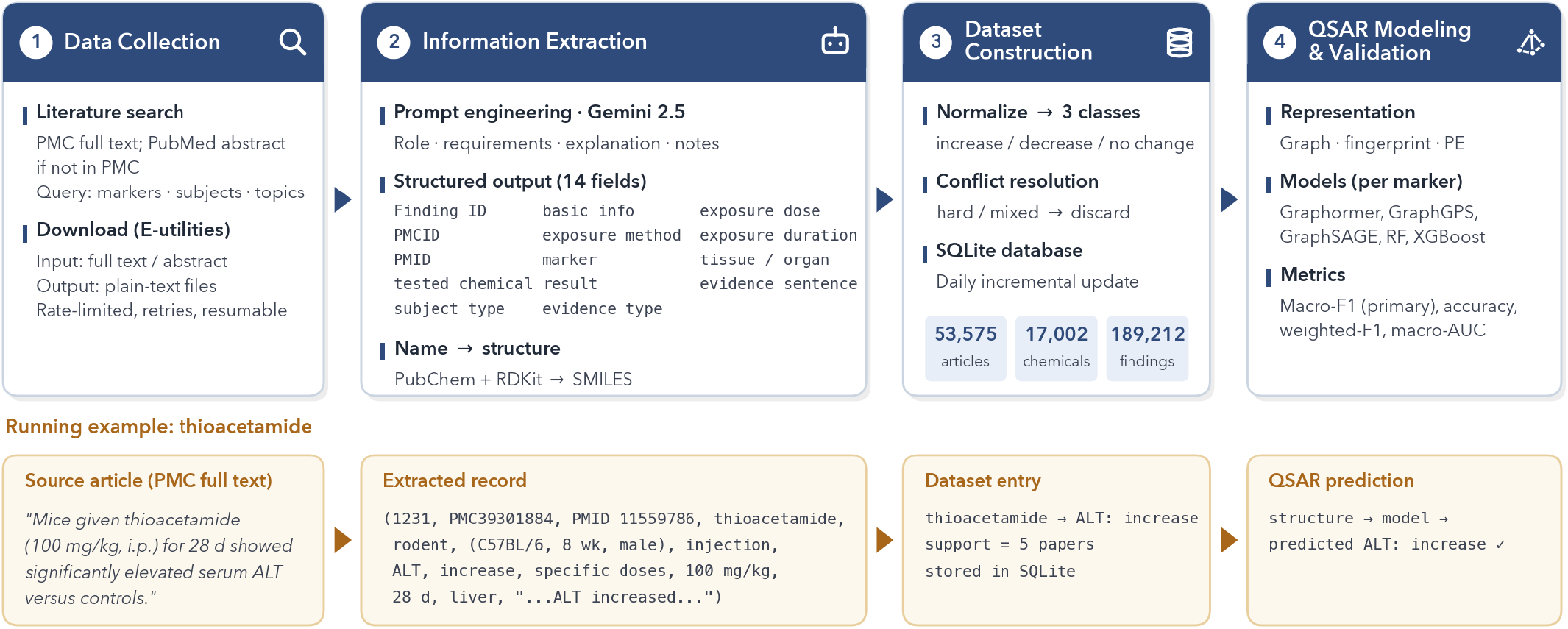
Overview of the proposed framework, which consists of four stages: data collection, information extraction, dataset construction, and QSAR baseline modeling.

### B. Data Acquisition

We search PubMed with a title/abstract (`[tiab]`) query that requires a match in each of three term groups: toxicity biomarkers (e.g., ALT, AST, ROS, MDA, ATP), test subjects or biological samples (e.g., cells, rats, mice, serum), and hepa-totoxicity topics (e.g., liver injury, DILI, steatosis, cholestasis). The query is limited to English-language journal articles and excludes studies on nanoparticles or microplastics as well as epidemiological, cohort, and survey studies, so that it focuses on experimental toxicology. The publication years range from 1995 to 2025. For each hit, we download the PMC full text when it is available and the PubMed abstract otherwise, using the NCBI E-utilities with rate limiting, retries, and a download registry that lets interrupted runs resume. The full term lists, the yearly article counts, and the download details are given in Appendix A.

The corpus reported here is a snapshot of the articles retrieved and processed between May 27 and June 11, 2026 (EDT); articles added by later daily updates are not included in the statistics. It contains 53,575 articles, 45,096 with full text and 8,479 with abstract only. Of these, 22,001 articles (≈41%) yielded extractable results, producing 189,212 findings across 17,002 chemicals.

### C. Dataset Construction

Records are kept in a single-file SQLite database. An *articles* table keyed by PMID holds article metadata, and a *findings* table links each finding to its source article through the PMID, so every finding can be traced back. Findings are written as soon as they are extracted, so an interrupted run does not lose processed results. A uniqueness constraint on `(PMID, tested_chemical, marker, result, evidence_sentences)`, combined with `INSERT OR REPLACE`, means that reprocessing an article overwrites identical records instead of duplicating them. The schema is shown in Appendix A (Fig. 4).

### D. Information Extraction

We use an LLM because the same result can be written in many ways (an *increase* may appear as “rise,” “elevate,” or “improve”), which rule-based methods, NER, and regular expressions handle poorly at the scale of our corpus.

#### a) Input and output

The input is the plain-text PMC full text or PubMed abstract, truncated to 50,000 characters. Truncation removes the end of a document, which usually holds the references and supplementary material, so findings in the main results sections are normally kept; findings reported only in late sections of very long articles may be missed. We did not count how many articles were truncated. We use Gemini 2.5 Flash (`gemini-2.5-flash`) [18]; all extraction runs reported in this paper were performed between May 27 and June 11, 2026 (EDT). The temperature is set to 0, which makes outputs as deterministic as possible but does not remove run-to-run variation. Each finding has eleven content fields (tested chemical, test subject, basic information, exposure route, biomarker, outcome, evidence type, exposure dose, exposure duration, tissue/organ, and evidence sentence) and three identifiers (Finding ID, PMCID, and PMID).

#### b) Prompt engineering

Prompt design strongly affects extraction accuracy [19] and hallucination [20]. Our prompt casts Gemini 2.5 as an expert in hepatotoxicity and has three parts. The first sets the output format and two constraints: single-valued fields with several values must be split into separate records, and if the four core elements (tested chemical, test subject, biomarker, and outcome) cannot all be found, the model returns only “No results.” The second gives rules for each field: inclusion and exclusion criteria (e.g., the tested chemical must be a single compound, not a class, mixture, solvent, or nanomaterial), controlled vocabularies, and the usual assays and synonyms of each biomarker. The third restates common failure cases: do not invent records, do not output markers outside the target list, and always copy a verbatim evidence sentence, which must contain the dose when a specific dose is reported. More detail is given in Appendix B.

### E. QSAR Modeling

#### a) Normalization and conflict resolution

The extraction prompt restricts findings to the effect of a single chemical tested alone, compared with a negative control group or assessed through a dose-dependent pattern; results from co-treatment or combination groups are explicitly excluded. By design, this leaves out the common protective-treatment setting, in which a compound is given together with an injury-inducing agent and compared with the injury model group. For each marker, we map the result field to three classes (increase, decrease, and no significant change) and discard entries that cannot be mapped. Findings are then aggregated per chemical, and a chemical is discarded if its reports are inconsistent. A *hard* conflict means that both “increase” and “decrease” were reported; a *mixed* conflict means that a directional result co-occurs with “no significant change.” A retained chemical takes the single result its papers agree on as its label, and its support is the number of distinct papers reporting that result. Disagreements are thus resolved by discarding, not by voting. Dose, species, and tissue are recorded for every finding but are pooled when these chemical-level labels are derived.

#### b) Name-to-structure conversion

Chemical names are converted to SMILES strings by querying PubChem [5], with lookups cached locally, and every SMILES is canonicalized with RDKit so that each compound has exactly one identifier [21]. Names that PubChem could not resolve to a structure (e.g., mixtures, extracts, and biological macromolecules) were excluded from QSAR modeling. No dedicated filter was applied to non-small-molecule agents such as lipopolysaccharide or concanavalin A; they were retained only if PubChem returned a structure for the name.

#### c) Models

We compare five models per marker. Random forest (400 trees, balanced class weights) and XGBoost (400 trees, maximum depth 6) use 2048-bit Morgan fingerprints with radius 2. Three GNNs (Graphormer, GraphGPS, and GraphSAGE) take molecular graphs with atoms as nodes and bonds as edges, and share a backbone of three graph layers (hidden size 128), global mean pooling, and an MLP classifier. Feature, architecture, and training details are given in Appendix C.

## IV. EXPERIMENTS

### A. Experimental Setup

We train one model per biomarker under three dataset settings. In the 3-class (FULL) setting, each chemical is labeled increase, decrease, or no significant change. The binary setting removes the no-significant-change class (increase vs. decrease). The high-confidence setting keeps only chemicals whose label is supported by at least two papers, since different articles may report different results for the same chemical– biomarker pair. Within each marker, chemicals are split at random (stratified by label) into training, validation, and test sets in a 64/16/20 ratio with a fixed seed (42). We did not perform hyperparameter search; all models use fixed settings shared across markers. We report macro-F1, accuracy, weighted-F1, and macro-AUC, together with per-class F1. Macro-F1 is the primary metric.

### B. Dataset Statistics

The most frequently studied chemicals are carbon tetra-chloride, acetaminophen, ethanol, thioacetamide, lipopolysac-charide, and concanavalin A. All are well-known experimental hepatotoxicants, and carbon tetrachloride, acetaminophen, thioacetamide, and lipopolysaccharide are standard agents for inducing liver injury in animal models [22]. The extraction therefore recovers the compounds expected in this domain (face validity). In a sample of 1,000 full-text PMC articles, findings concentrate on a small set of biomarkers (Fig. 2): liver-function markers (ALT, AST, ALP, bilirubin) and histopathological endpoints (steatosis, fibrosis) are reported most often, and most other markers form a long tail. A breakdown of the same sample by subject, marker, and result is shown in Appendix A (Fig. 5).

**Fig. 2.**
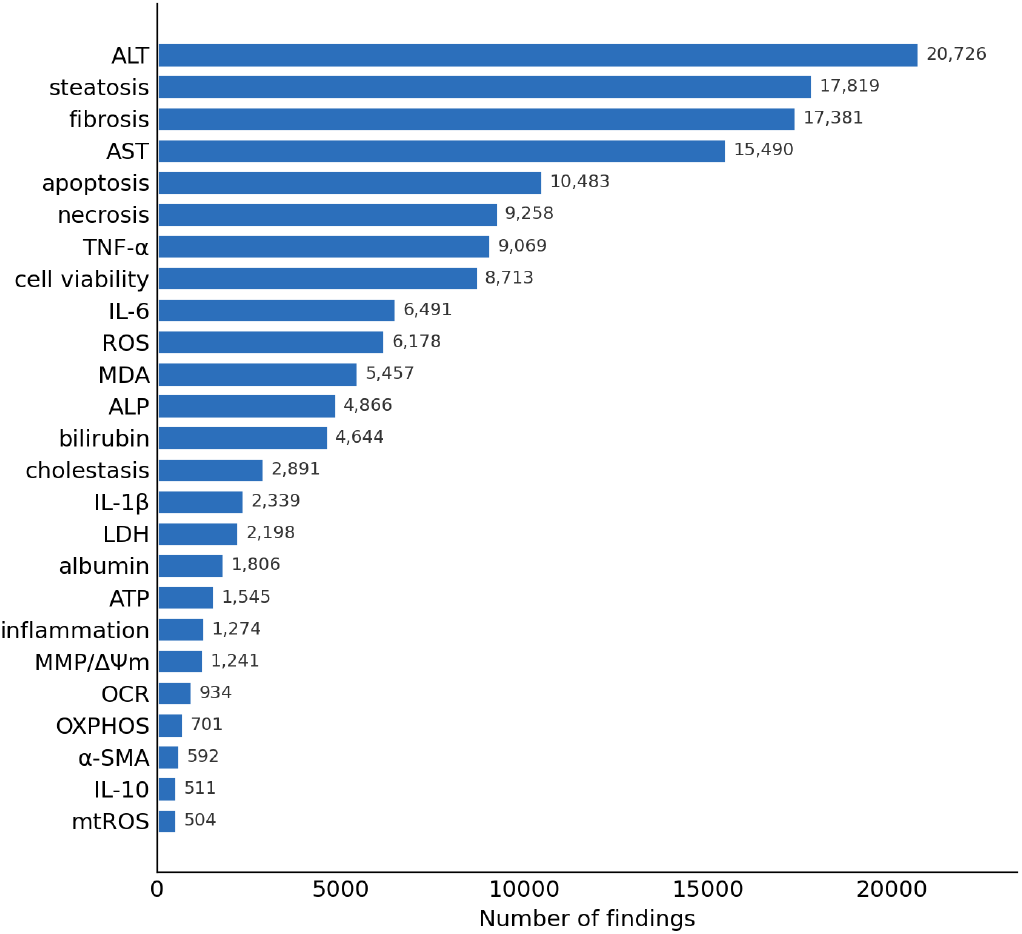
Distribution of findings across the 25 most frequent biomarkers based on 1,000 full-text papers. Liver-function markers (ALT, AST, ALP, bilirubin) and histopathological endpoints (steatosis, fibrosis) are the most frequently reported, while the majority of markers form a long tail.

**Fig. 3.**
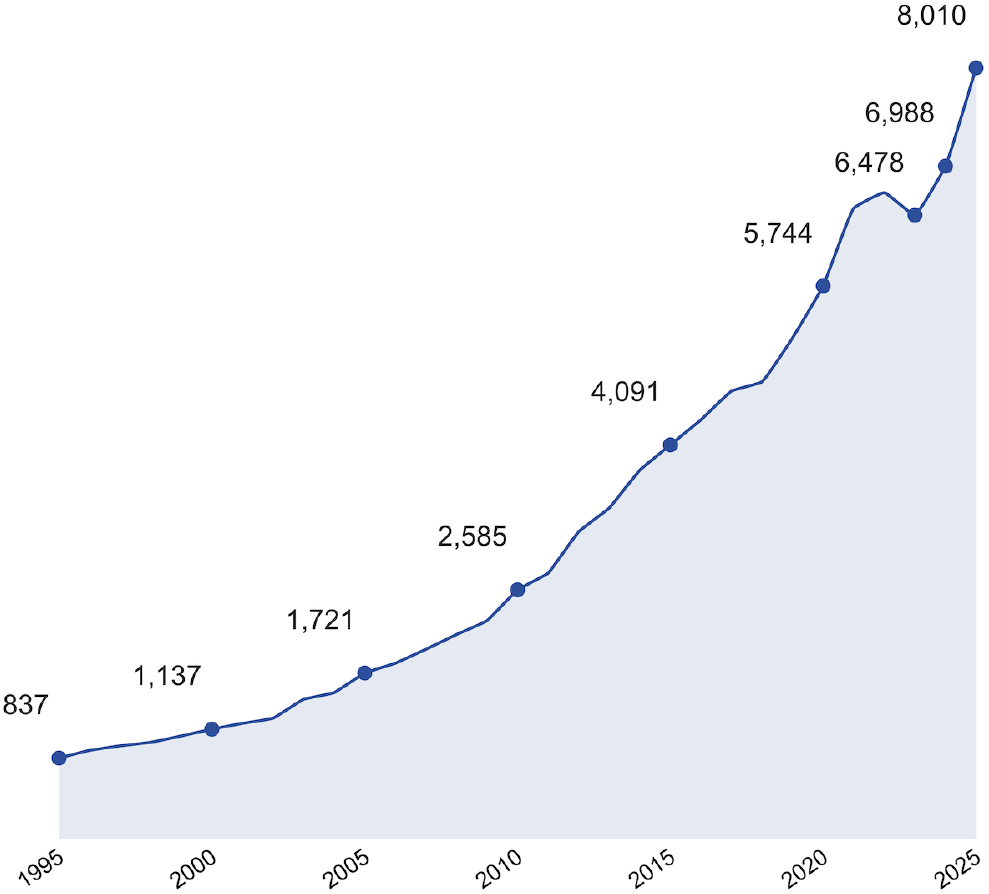
Annual number of retrieved articles in the corpus (1995–2025). The yearly count rises steadily from 837 in 1995 to 8,010 in 2025, reflecting the growing body of hepatotoxicity literature over the past three decades. The number of articles published in 2026 is not included in this figure.

**Fig. 4.**
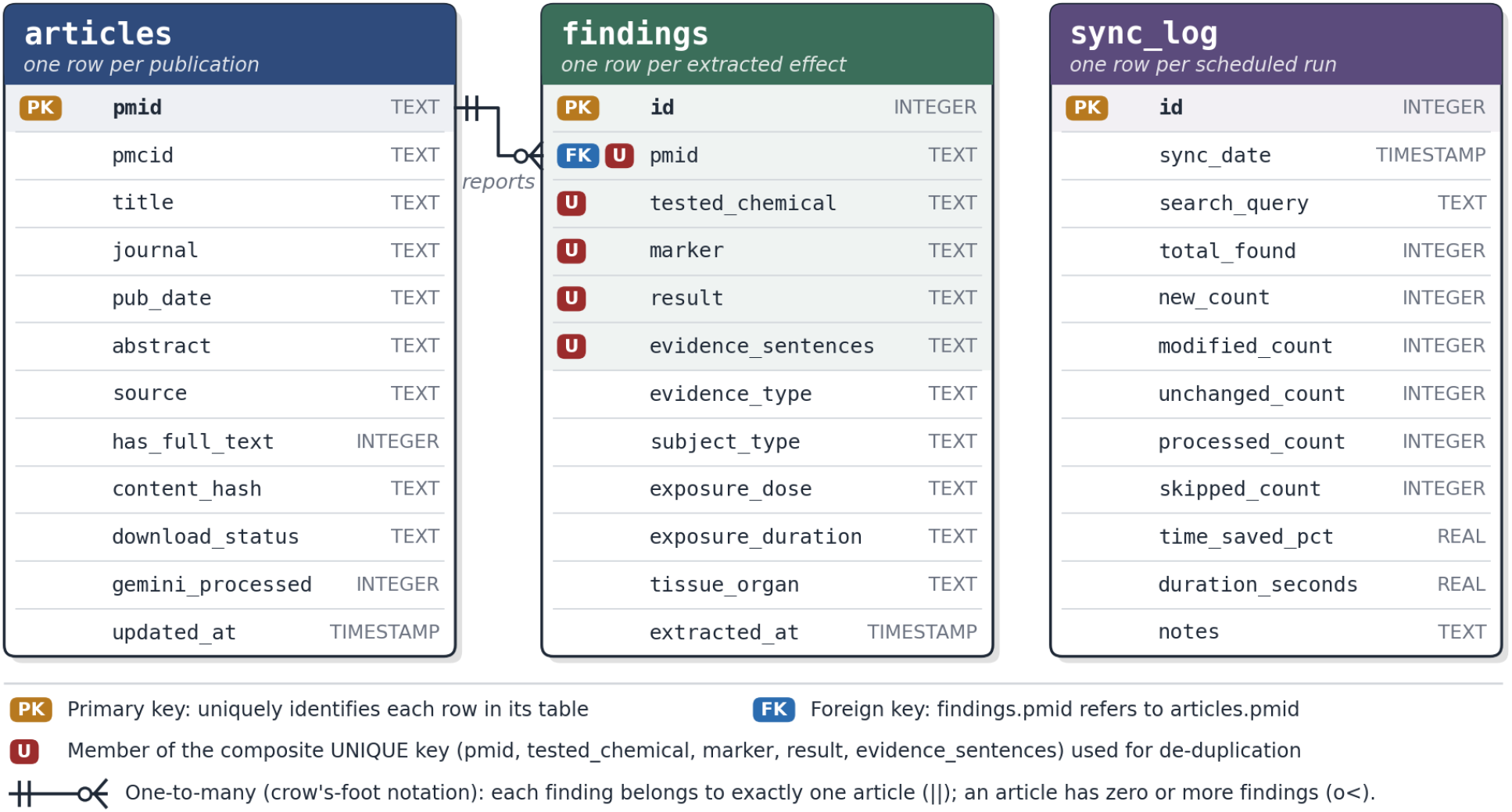
Entity–relationship diagram of the incrementally updated SQLite database. `articles` stores one row per publication, keyed by PMID, together with bibliographic metadata and per-article pipeline state (e.g., the content hash used for change detection). `findings` stores one row per extracted effect and references its source article through PMID (one-to-many); findings are de-duplicated by a composite uniqueness constraint (U). `sync_log` records one row per scheduled run. Selected columns are shown.

**Fig. 5.**
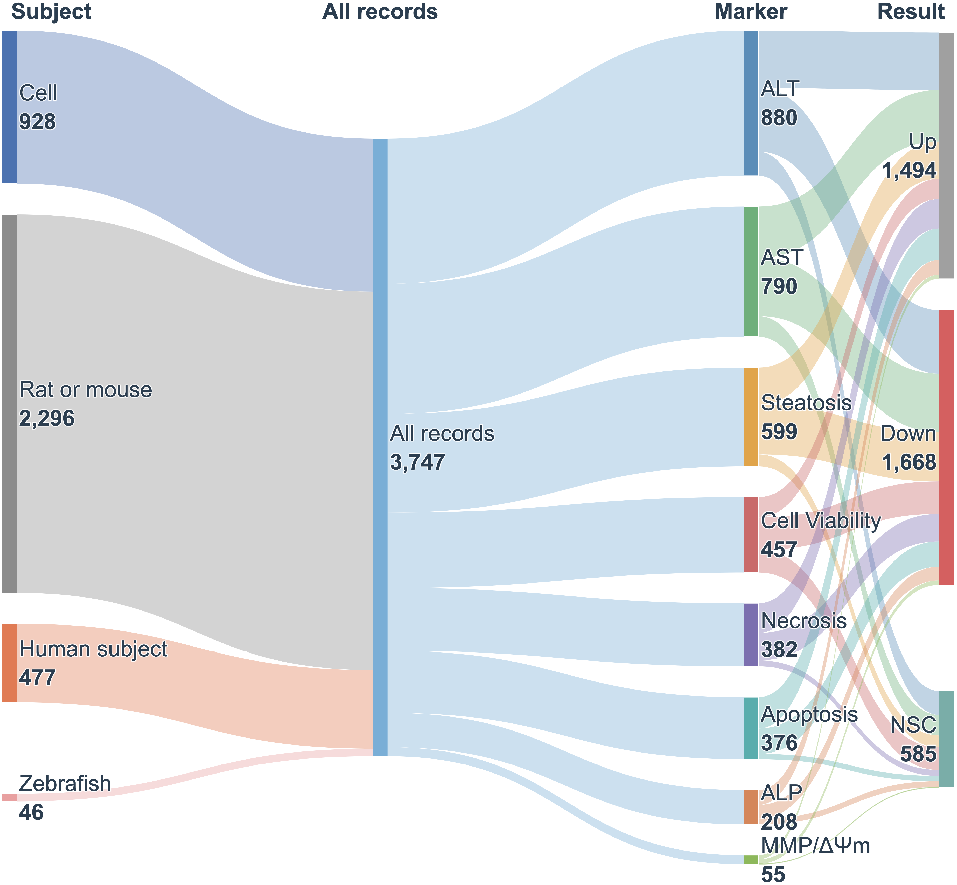
Sankey diagram showing the composition of the records obtained by text mining from a sample of 1,000 full-text PMC articles (3,747 records in total). The most frequently reported marker was ALT (23%), followed by AST (21%) and steatosis (16%). The proportion of records reporting a decrease in the measured marker (45%) was slightly higher than those reporting an increase (40%), while few records reported “no significant change” (NSC, 16%). In terms of experimental subjects, rats or mice, cells, human subjects, and zebrafish accounted for 61%, 25%, 13%, and 1%, respectively.

### C. Extraction Quality

To assess extraction quality, we randomly sampled 35 full-text articles published in 2025, which yielded 217 extracted findings. A domain expert in hepatotoxicology read each source article in full and, for every extracted finding, determined the correct field values from the article text. Each finding was compared field by field with this ground truth and labeled as *fully correct* (all fields agree with the source), *incomplete* (the core chemical–marker–result triple is correct, but a secondary field, mainly the exposure dose, is missing or not traceable to the evidence sentence), or *incorrect* (at least one element of the core triple disagrees with the source, or the record should not have been extracted). Of the 217 findings, 208 (95.9%) were fully correct, none was incomplete, and the remaining 9, all from 2 articles, were incorrect.

### D. QSAR Results Across Markers

Of the 235 markers, 16 have at least 500 labeled chemicals and 74 have more than 50. Because sample sizes differ widely, Table I reports the 16 markers with *N* ≥ 500; results for the remaining 58 markers are in the repository and are marked as small-sample, high-variance results.

**TABLE 1.**
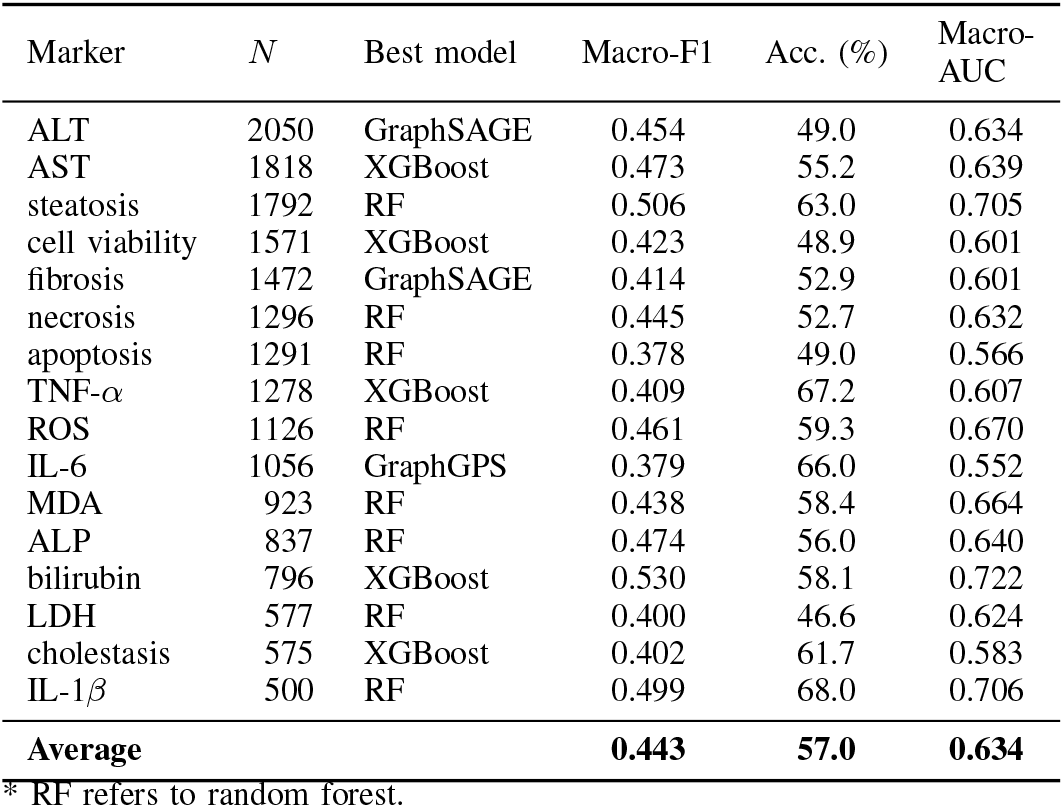
Per-markeR QSAR RESULTS ON THE 16 RELIABLE MARKERS (*N* ≥ 500 CHEMICALS), 3-CLASS SETTING, SORTED BY *N*. *N* IS THE NUMBER OF CHEMICALS; FOR EACH MARKER, THE MODEL WITH THE HIGHEST TEST MACRO-F1 AMONG THE FIVE MODELS IS REPORTED. BECAUSE THIS SELECTION USES THE TEST SET, THESE VALUES ARE OPTIMISTIC AND SHOULD BE READ AS UPPER BOUNDS.

| Marker | *N* | Best model | Macro-F1 | Acc. (%) | Macro-AUC |
|---|---|---|---|---|---|
| ALT | 2050 | GraphSAGE | 0.454 | 49.0 | 0.634 |
| AST | 1818 | XGBoost | 0.473 | 55.2 | 0.639 |
| steatosis | 1792 | RF | 0.506 | 63.0 | 0.705 |
| cell viability | 1571 | XGBoost | 0.423 | 48.9 | 0.601 |
| fibrosis | 1472 | GraphSAGE | 0.414 | 52.9 | 0.601 |
| necrosis | 1296 | RF | 0.445 | 52.7 | 0.632 |
| apoptosis | 1291 | RF | 0.378 | 49.0 | 0.566 |
| TNF-*α* | 1278 | XGBoost | 0.409 | 67.2 | 0.607 |
| ROS | 1126 | RF | 0.461 | 59.3 | 0.670 |
| IL-6 | 1056 | GraphGPS | 0.379 | 66.0 | 0.552 |
| MDA | 923 | RF | 0.438 | 58.4 | 0.664 |
| ALP | 837 | RF | 0.474 | 56.0 | 0.640 |
| bilirubin | 796 | XGBoost | 0.530 | 58.1 | 0.722 |
| LDH | 577 | RF | 0.400 | 46.6 | 0.624 |
| cholestasis | 575 | XGBoost | 0.402 | 61.7 | 0.583 |
| IL-1*β* | 500 | RF | 0.499 | 68.0 | 0.706 |
| **Average** |  |  | **0.443** | **57.0** | **0.634** |

\* RF refers to random forest.

We report these results as baseline benchmarks. On the 16 markers, the average macro-F1 is 0.443, accuracy is 57.0%, and macro-AUC is 0.634, all above the three-class random baselines (macro-F1*≈* 0.33, AUC = 0.5). Structure alone thus predicts a chemical’s effect direction on a marker better than chance. The moderate level of performance is also consistent with label noise in the extracted data and with the limited predictability of these endpoints from structure, so these results should be read as baselines for future work rather than as a validation of the dataset.

Finding 1: at the current data scale, the classic models remain competitive. Counting the best model per marker, the classic models (random forest and XGBoost) win on 44 of the 74 markers and the GNNs on 30. On the 16 data-rich markers the gap is larger: the classic models win 13 (random forest 8, XGBoost 5) and the GNNs only 3. Because no hyperparameter tuning was done for any model and no significance t est w as r un, t his c omparison i s only indicative. It is, however, in line with reports that on real-world literature data with limited samples and noisy labels, strong fingerprint-based b aselines a re o ften m ore r obust than more heavily parameterized GNNs [23]. Finding 2: performance varies across markers. The best predictions are obtained for bilirubin (macro-F1 0.530), steatosis (0.506), and IL-1*β* (0.499), whose effect directions are relatively unambiguous. In contrast, apoptosis (0.378) and IL-6 (0.379) are close to random; their reported effect directions are more dispersed and harder to infer from structure alone. Finding 3: small-sample markers. Occasional high scores among the remaining 58 markers (e.g., TBIL, PPAR*α*) mainly reflect t he variance of very small test sets. We take them only as evidence of the breadth of coverage and do not interpret them individually.

## V. CONCLUSION

We presented and evaluated an LLM-based pipeline that mines hepatotoxicity findings from the literature. From 53,575 PubMed and PMC articles, Gemini 2.5 extracted 189,212 structured findings spanning 235 markers, stored in a SQLite database that is updated daily. In a manual check, 95.9% of the sampled findings were fully correct; the 9 incorrect findings came from only 2 articles and followed recurring patterns, which suggests systematic rather than random errors, although a larger evaluation set is needed to confirm this. Per-marker QSAR models reached an average macro-F1 of 0.443 on the 16 data-rich markers, above the random baselines, and classic models remained competitive with GNNs. Future work will improve entity normalization and disambiguation, add combined-exposure data, use cross-validation, and extend the pipeline to other endpoints such as nephrotoxicity and neurotoxicity.

## DATA AND SOFTWARE AVAILABILITY

All source code, the constructed dataset, the full extraction prompt, the complete search query, and the manual-validation data used in this work are available at the following repository: https://github.com/jiqiu-win/biolit_autominer

## ACKNOWLEDGMENT

Chunjiang Zhu was partially supported by Natural Science Foundation CCF-2609947, 4-VA grant 183643, and Old Dominion University Startup Funds.

## APPENDIX A

### DATA ACQUISITION AND DATABASE DETAILS

#### a) Search query

Candidate articles are retrieved with a PubMed query restricted to the title/abstract field (`[tiab]`). An article must match all three of the following term groups (terms within a group are combined with OR, and the three groups are combined with AND):

- **(i) Toxicity biomarker terms**, covering liver-function markers (e.g., ALT, AST, ALP, bilirubin, albumin, LDH) and mitochondrial/toxicity-related markers (e.g., ROS, MDA, ATP, mitochondrial membrane potential/MMP, mtROS, OCR, mPTP, cytochrome c release, as well as TNF-*α*, IL-6, apoptosis, necrosis, and mitochondrial dysfunction/damage);
- **(ii) Subject or biological-sample terms**: cells, rats, mice, zebrafish, and biological samples such as serum, blood, plasma, and urine;
- **(iii) Hepatotoxicity topic terms**, such as hepatotoxicity, liver/hepatic toxicity, liver injury/damage, drug-induced liver injury (DILI), hepatocyte injury/necrosis/apoptosis, steatosis, cholestasis, and liver fibrosis.

The query is further limited to journal articles (`“journal article”[pt]`) and English-language articles (`English[la]`), and a `NOT` clause excludes articles focused on nanoparticles or microplastics, as well as epidemiological, cohort, and survey studies. Fig. 3 shows the yearly number of retrieved articles.

#### b) Download pipeline and robustness

We use the of-ficial E-utilities [24] to run the query and download the literature. We first obtain the list of PMIDs matching the query via `esearch`, and then fetch each article via `efetch` and the PMC Open Access interface, falling back to the PubMed abstract when full text is not available in PMC. To comply with NCBI’s access policy, we cap the request rate at its documented limits (about 10 requests/s with an API key and about 3 requests/s otherwise) and set a timeout on every network request. Each failed request is retried up to three times with an increasing back-off; if it still fails, the item is logged and skipped without interrupting the run. Transient server-side errors from PMC (e.g., 502/503/504 and connection timeouts) are distinguished from failures on our side and deferred to a later retry. A download registry skips already-downloaded articles, which avoids redundant downloads and lets an in-terrupted run continue. Each downloaded file is checked to confirm that it is genuine full text (non-empty, at least 150 words, and free of the “abstract-only” placeholder marker); files that fail the check are flagged or fall back to the abstract. The daily incremental updates reuse the same pipeline.

#### c) Database schema

All records are stored in a SQLite database, a lightweight, serverless relational database kept in a single file (Fig. 4). The *articles* table, keyed by PMID, holds one row per article with its metadata (title, journal, publication date, and PMCID). The *findings* table, keyed by an auto-increment id, holds the extracted fields and links to its source article through the PMID. A *sync log* table records each incremental update, and a key–value *meta* table stores dataset metadata. Findings are written row by row at extraction time rather than bulk-loaded at the end. The *articles* table is unique by PMID, and a write busy-timeout makes concurrent writes wait and be serialized safely.

## APPENDIX B

### PROMPT DESIGN DETAILS

The prompt is organized into three parts: extraction requirements, element-by-element specifications, and important reminders.

The first part prescribes a structured output and establishes two key constraints: (i) the elements within a single record must be mutually consistent—seven fields are single-valued, and multiple values must be split into separate records, whereas dose, duration, and tissue/organ may take multiple values; and (ii) tested chemical, test subject, biomarker, and outcome are the four core elements, and if they cannot be extracted jointly the model returns only “No results,” with uninformative fields filled by “-”.

The second part provides precise rules for each element:(1) exhaustive inclusion and exclusion criteria (e.g., the tested chemical must be a single, specific compound, excluding chemical classes, mixtures, environmental matrices, solvents, microplastics, and nanomaterials); (2) controlled vocabularies for several elements (test subject drawn from six categories; outcome restricted to increase/decrease/no significant change; evidence type restricted to `specific doses`); (3) fixed output formats; (4) explicit logical dependencies among elements; and (5) illustrative terminology with explanations of key abbreviations and units and, for biomarkers, the common assays and synonymous spellings of each indicator (e.g., ROS mapped to DCFH-DA and its aliases).

The third part emphasizes error-prone cases. For instance, “do not fabricate any record; output No results when no information is present” targets hallucination; “extract only the designated biomarkers (do not output SOD, GSH, CAT, etc.)” curbs out-of-scope extraction; and “every record must include an evidence sentence copied verbatim from the source text, and when the evidence type is `specific doses` the evidence sentence must contain the actual dose value” gives each record verifiable provenance. The full prompt is available in the repository.

## APPENDIX C

### QSAR MODEL DETAILS

#### a) Molecular representation

For the classic models, each molecule is encoded as a 2048-bit Morgan (extended-connectivity) fingerprint with radius 2, where each bit indicates the presence of a particular local substructure. For the graph-based models, each molecule is an attributed graph with atoms as nodes and bonds as edges. Every atom is described by a 27-dimensional feature vector encoding its element type. Every bond is described by a 6-dimensional feature vector encoding its type (single, double, triple, or aromatic), conjugation, and ring membership, and is represented as two directed edges so that information propagates in both directions during message passing. For GraphGPS, each node additionally receives a Laplacian positional encoding, defined as the first eight non-trivial eigenvectors of the symmetric normalized graph Laplacian.

#### b) Models

The random forest uses 400 trees with balanced class weights. The XGBoost classifier uses 400 trees, maximum depth 6, learning rate 0.1, and subsample and column-subsample of 0.8. The graph models share a common backbone: the 27-dimensional atom features are linearly projected to a hidden size of 128, updated by three graph layers, aggregated by global mean pooling, and classified by a two-layer MLP head, with a dropout of 0.3 throughout. They differ in how each layer updates node representations. **Graphormer** stacks graph-transformer layers, each applying multi-head self-attention followed by a feed-forward sublayer, with residual connections and layer normalization; node representations are further augmented with a degree embedding. **GraphGPS** stacks GPS layers, each combining a local message-passing convolution with global multi-head self-attention, and uses the 8-dimensional Laplacian positional encoding. **GraphSAGE** stacks SAGE convolutions with residual connections and layer normalization; it uses only node features and connectivity, ignoring bond features, and serves as a simpler message-passing baseline.

#### c) Training

The graph models are trained with Adam (learning rate 10^−3^, batch size 64) and a class-weighted cross-entropy loss for up to 100 epochs, with early stopping (patience 20) on validation macro-F1; the checkpoint with the best validation macro-F1 is evaluated on the test set. All experiments use a fixed random seed (42). The GNNs are trained on a single GPU with PyTorch and PyTorch-Geometric, and the classic baselines use scikit-learn and XGBoost. SMILES canonicalization, molecular-graph construction, and Morgan fingerprints are computed with RDKit.

## APPENDIX D

### DISCUSSION

#### A. Limitations

This study has several limitations. **First, combined exposures and mixtures were excluded**. So that each record refers to one well-defined substance, we excluded chemical classes, mixtures, and nanomaterials. Real-world toxicity often comes from several substances acting together, and the current dataset cannot describe such exposure scenarios. **Second, the choice of classification granularity involves trade-offs**. We mapped effects to “increase,” “decrease,” and “no significant change.” The last category has very few samples for many biomarkers, which makes the three-class task hard to learn for data-sparse markers. Binary classification (increase vs. decrease) improved accuracy but loses the ability to report “no effect,” which is itself an important conclusion. Neither option is best in every case. **Third, no comparison across LLMs or prompts**. All extractions used a single LLM (Gemini 2.5) and a single prompt design. We did not compare other LLMs or prompting strategies, so the reported extraction quality is specific to this configuration and may not carry over to other models. **Finally, evaluation noise arises from a single random data split**. QSAR results come from one random, label-stratified train/validation/test split with a fixed seed. Random splits can place structurally similar compounds in both training and test sets and may therefore overestimate performance compared with scaffold-based splits, and a single seed gives no estimate of variance. Evaluation with scaffold splits and multiple seeds is left for future work.

#### B. Implications and Outlook

The dataset can serve as a watchlist of suspected environmental pollutants and drug candidates for hazard screening, and as training and benchmark data for QSAR, GNN, and read-across studies. Its chemical–marker–effect triples also allow analyses of substructure–effect associations and of research hotspots and gaps. Because the pipeline runs daily, the resource grows with the literature instead of staying a one-time snapshot.

## Notes

### Competing Interest Statement

The authors have declared no competing interest.

